# Large-scale production of melanin nanoparticle from *Pseudomonas stutzeri* strain BTCZ109

**DOI:** 10.64898/2026.07.10.737634

**Authors:** Dayana Mathew, Sarita G. Bhat, M. Chandrasekaran

## Abstract

Past few decades witnessed a boom in pharmaceutical and bioproduct industry with the help of bioprocess technology. Industrially important bioproducts can be produced in large scale for commercialization with the help of fermenters. Here in, pharmaceutically valuable bioproduct melanin, synthesized from *Pseudomonas stutzeri* strain BTC109 by using two different sized bioreactors. Under controlled conditions the bacteria were allowed to synthesis melanin nanoparticles. The important parameters to be monitored here are pH, dissolved oxygen, agitation, aeration, melanin production and cell biomass concentration.

**Graphical abstract:** 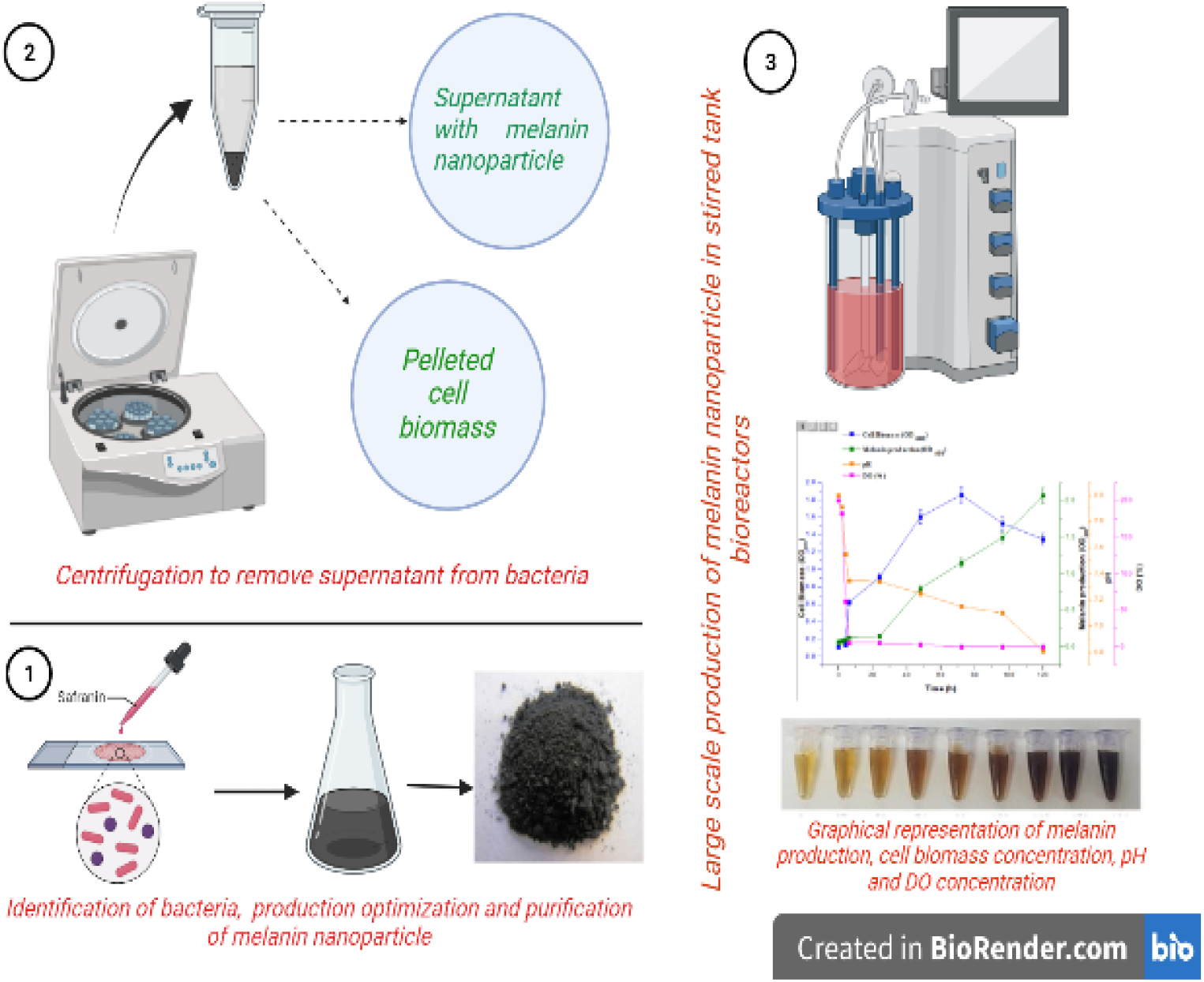

**Highlights:**

- Pharmaceutical and bioproduct development industries witnessed a shoot up due to bioprocess technology.
- Industrially important bioproduct like melanin can be produced in large scale with the help of industrial fermentation technology.
- The product thus obtained was found to be nano sized and it can be commercialized.

## 1. Introduction

Melanin are natural biopolymers synthesized by all forms of life starting from unicellular prokaryotes to advanced eukaryotes. These are polyphenolic heteropolymers synthesized mainly by classical Mason-Raper biosynthetic pathway (Ammanagi *et al*., 2019). There are several kinds of melanin found in living organism like, allomelanin, neuromelanin, pheomelanin, eumelanin, pyomelanin etc (Cao *et al*., 2021; Pralea *et al*., 2019) are some among them. The biological properties of this molecule made it industrially more valuable. The important properties of melanin including its UV protection ability, radical scavenging activity, metal chelation ability, antibiofilm effect, anticancer property, liver protection ability etc (Roy *et al*., 2022; Liu *et al*., 2022; Guo *et al*., 2023) enhanced the demand of the molecule commercially. Melanin also found to have nano structural properties which increases the biomedical properties of the molecule. As a nanoparticle it can be used as drug delivery vehicle, bioimaging agent, bioelectronics etc (Park *et al*., 2019; Michael *et al*., 2023).

The manufacturing of pharmaceuticals and bioproducts in large scale bioreactors witnessed significant advances with the help of both mammalian and microbial cell system for the past few decades. Various industrially important bioproducts like, antibiotics (Sánchez *et al*., 2010), pharmaceutical proteins (Nielsen *et al*., 2013), industrial enzymes (Niyonzima *et al*., 2019), organic acids (Sauer *et al*., 2008) etc., have been synthesized by utilizing industrial fermentation. More than 300 million USD were utilized in worldwide market for production of biotechnological products (Neubauer., 2011). Compared to chemical reaction processes, industrial fermentation is more useful because it incorporates natural resources under moderate cultural conditions involving both pressure and temperature for production of bioproducts with considerably increased interest and promoted emergence in bio-economy (Milne, N., 2020; Sáez-Sáez *et al*., 2020). Engineered cell factories were obtained with the involvement of genetic and metabolic engineering along with synthetic biology which are capable of producing improved products with enhanced product yield. The process development is always at a lag from 3-10 years with increased cost of up to 100-1 million USD.

Here in this study, large scale production of melanin nanoparticles was carried out from bacterial strain *Pseudomonas stutzeri* BTCZ109. Two different sized fermenters of 2L and 5L capacities were used for this purpose. The important media components optimized through one-factor-at-a-time method and statistical design of experiments were used for media design. In the fermenter the components pH, temperature, aeration, agitation etc. were maintained in a controlled condition for large scale production of melanin nanoparticles. The important components like melanin production concentration, cell biomass, pH and dissolved oxygen were monitored at regular intervals of sampling. After production and purification, the molecule was evaluated to identify the nano structural property through several characterization methods.

## 2. Materials and method

### 2.1 Bacterial strains for melanin production

The microorganism, *Pseudomonas stutzeri* BTCZ109, used in this study, was maintained at 20 °C on Yabuuchi and Ohyama’s tyrosine basal agar slants with 2g/L L-tyrosine (Yabuuchi, & Ohyama, 1972) (Composition of the media: sodium chloride (NaCl) 5g/L, magnesium sulfate (MgSO_4._ 5H_2_O) 0.1g/L, potassium dihydrogen phosphate (KH_2_PO_4_.7H_2_O) 2g/L at a pH of 8) and transferred monthly.

### 2.3 Large scale production of melanin nanoparticle using bioreactor

By using bioreactors industrial production of biopolymer like melanin’s are possible by eliminating several restrictions encountered in shake flask cultures. The operation conditions should be optimized for maximum production of melanin nanoparticles. Large scale production of this biopolymer is made possible by using batch mode of fermentation.

### 2.4 Fermentation carried out in different sized fermenters

#### 2.4.1 Fermentation at 2 L Scale

Mechanically agitated stirred tank bioreactors are used in lab scale fermentation experiment with a total capacity of 2L and a working volume of 1L, with height to diameter ration of 1.0 with one impeller with four bladed turbines mounted on vertical shaft. The inoculum was first prepared by transferring a loop full of inoculum to 5mL nutrient broth in a test tube. It was kept at rotary shaker at 120 rpm for 37 ^0^C overnight. Inoculum preparation was carried out in 500 mL Erlenmeyer flask with a volume of 250 mL medium of nutrient broth and sterilized prior to inoculation. From the overnight bacterial culture 200μL of inoculum was transferred to freshly prepared nutrient media. Flasks were then incubated at rotary shaker at 120 rpm at 37 ^0^C for 24 h incubation. To a working volume of 1L fermenter 8% (v/v) of inoculum was inoculated.

The fermenter parameters like pH, temperature, dissolved oxygen and agitation speed were monitored. The main fermenter operating conditions were kept at temperature 37 ^0^C, aeration 0.5-1 l/p/m, agitation speed 140 rpm (2.3 Hz) and pH maintained at 8.0 using 2.0 mol of 1L NaOH and 2.0 mol of 1L HCl. The fermentation process was monitored with sampling at 12h intervals. The samples collected were tested for sterility, pH, biomass content and melanin production concentration.

#### 2.4.2 Fermentation at 5L Scale

Mechanically stirred tank fermenters with 5L capacity with a working volume of 4L, a ratio of 1.5 height-to diameter and two vertically mount impellers with six-bladed turbines on a shaft was used for fermentation purpose. Water jacket connected externally in the fermenter vessel for maintaining the temperature. 4L of fermentation medium composed of sodium chloride (NaCl) 5g/L, magnesium sulfate (MgSO_4._5H_2_O) 0.1g/L, potassium dihydrogen phosphate (KH_2_PO_4_.7H_2_O) 2g/L at a pH of 8 was used for the purpose. The fermentation media was sterilized at 121 ^0^C for 15-20 min before inoculation. The inoculum was first prepared by transferring a loop full of inoculum to 5mL nutrient broth in a test tube. It was kept at rotary shaker at 120 rpm for 37 ^0^C overnight. From the overnight bacterial culture 200μL of inoculum was transferred to freshly prepared 500 mL nutrient media in a 1000mL flask. This was then kept in rotary shaker at 120 rpm for 24 h at 37 ^0^C in shake flasks. After 24 h the inoculum was used to inoculate the fermenter vessel with 5L capacity and working volume of 4L. The different parameters like pH, temperature, dissolved oxygen and agitation were monitored. The main fermenter operating conditions were kept at temperature 37 ^0^C, aeration 0.5-1 l/p/m, agitation speed 140 rpm (2.3 Hz) and pH maintained at 8.0 using 2.0 mol of 1L NaOH and 2.0 mol of 1L HCl. The fermentation process was monitored with sampling at 12h intervals. The samples collected were tested for sterility, pH, biomass content and melanin production concentration.

#### 2.4.3 Analytical methods

After 12 h of incubation samples were taken out from the fermenters and split in to two aliquots. The aliquoted samples were then measured for determining the maximum melanin concentration and cell biomass yield. One of the aliquots was centrifuged at 10,000 rpm for 15 min and the OD was checked at 400 rpm for checking melanin production. The other sample before centrifugation was analyzed at OD 600 to determine cell biomass concentration.

### 2.5 Downstream recovery process

Extraction of melanin nano particle was done by using 1 N HCl for 1 week of incubation and the melanoids were removed by heating. The cell debris were removed by centrifugation at 10,000 rpm for 10 min. Purity check was performed by using thin layer chromatogram using silica gel plates.

### 2.6 Melanin as nanoparticle

The nano structural distribution of melanin nanoparticle was analyzed with the help of different characterization methods involving UV spectral analysis, diffractive light scattering (DLS), zeta potential distribution, Transmission electron microscopy (TEM) etc.

#### 2.6.1 UV visible spectroscopy

The UV-Visible spectrum (Shimadzu UV-1800) of bacterial melanin was prepared (1mg/mL) in 0.1N NaOH and UV-Visible absorption spectrum was measured between 200-800 nm using 0.1 N NaOH as blank and compared.

#### 2.6.2 Dynamic light scattering

Dynamic Light Scattering size distribution and zeta potential or surface charge were determined (Zeta sizer Nano ZS, Malvern) at 25°C. Three independent measurements were performed.

#### 2.6.3 Transmission electron microscope

For TEM imaging, the nanogranules were dispersed in double distilled water, sonicated as above and separated from the mixture by centrifugation at 18,000 rpm for 15 min. The samples were placed on copper grids and air-dried before imaging using a JEOL JEM-1210 electron microscope at an operating voltage of 200 kV with a LaB6 source. The TEM images were taken before and after sonication to determine the exact size of this particle.

## 3. Results

### 3.1 Single batch production

Batch mode of fermentation is most desired in the case of melanin production, with one time addition of all desired nutrients into the fermenter. L-tyrosine which served as sole source of carbon and nitrogen was used in batch fermentation process. The only restriction with batch mode of fermentation is that nutrients are provided initially without any intermediate addition which restrict the bacterial growth on depletion of nutrients. Melanin is secondary metabolite, produced at the stationary phase of bacterial growth.

### 3.2 Fermentation at 2 L Scale

The temperature and pH were controlled at 37^0^C and 8.0, respectively in lab scale batch fermentation process. The initial media composition consists of sodium chloride (NaCl) 1.5 g/L, magnesium sulfate (MgSO_4._ 5H_2_O) 0.2 mM, L-tyrosine 6 g/L, potassium dihydrogen phosphate (KH_2_PO_4_.7H_2_O) 1.5g/L was used for the purpose.

Melanin production and cell biomass were monitored by sampling at 12 h interval for about 168 h. The maximum melanin production and cell biomass was obtained at 168h of incubation. The process of melanin production was depicted clearly in Figure 1. using 2L fermenter vessel. The cell biomass reached the maximum OD (Xmax) of 0.607 wet weight at 168 h, while the melanin concentration reached the maximum (Pmax) of 388.35 μg/mL at 168 h. The overall melanin formation rate (Qp) of 0.733 μg/mL l68 h.

**Figure 1.**
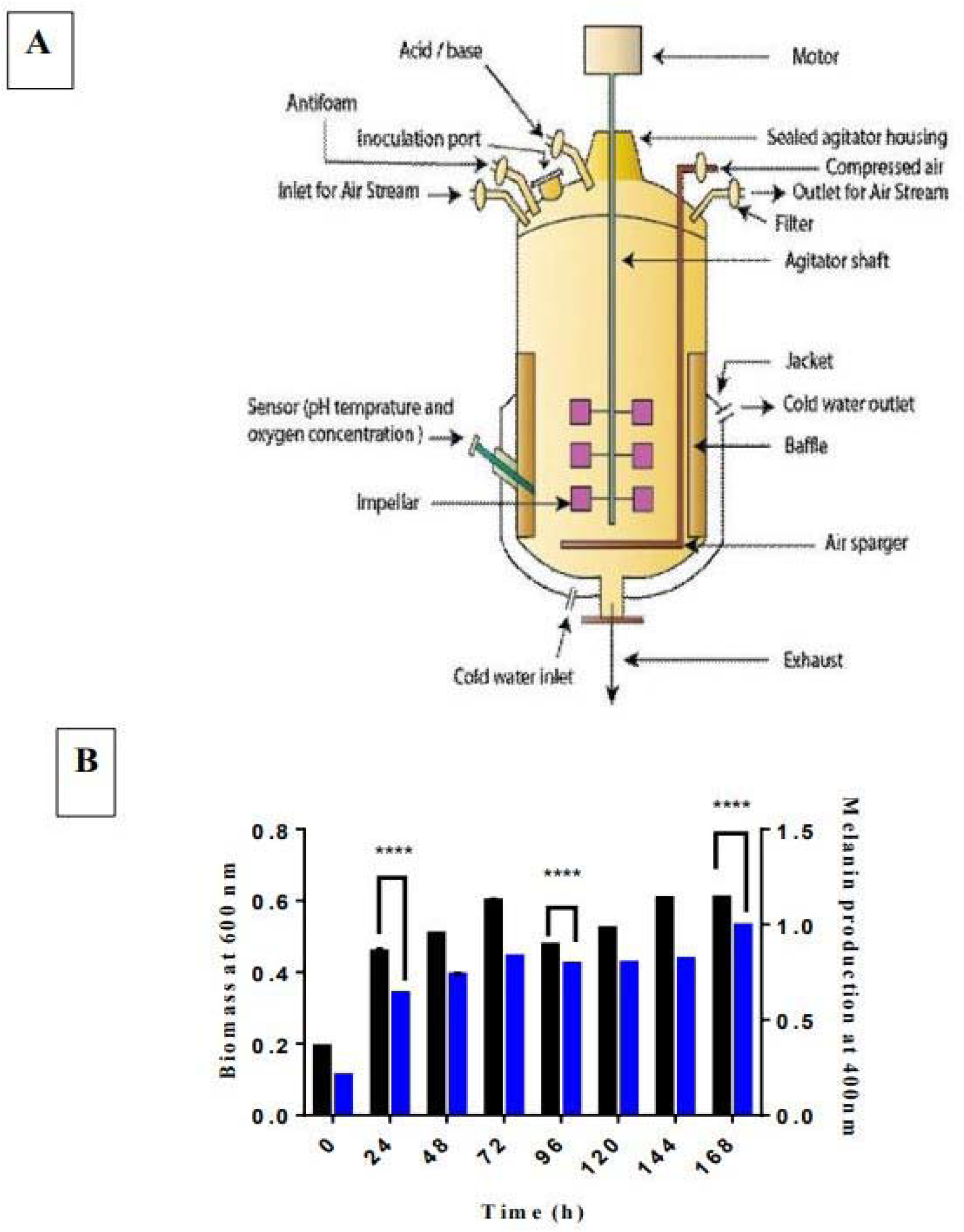
(a) Pictorial representation of fermenter (b) Time dependent production of biomass and melanin in 2L fermenter.

### 3.3 Fermentation at 5 L Scale

The temperature and pH were controlled at 37^0^C and 8.0, respectively in lab scale batch fermentation process. The initial media composition consists of sodium chloride (NaCl) 1.5 g/L, magnesium sulfate (MgSO_4._ 5H_2_O) 0.2 mM, L-tyrosine 6 g/L, potassium dihydrogen phosphate (KH_2_PO_4_.7H_2_O) 1.5g/L was used for the purpose. The dissolved oxygen (DO) concentration was considered at 100% and reduction in percentage of oxygen was monitored in the control panel.

Melanin production and cell biomass were monitored by sampling at 12 h interval for about 168 h. Consistent increase in melanin production was observed with respect to increase in time. The maximum melanin production and cell biomass was obtained at 120 h of incubation. The process of melanin production was depicted clearly in Figure 2. using 5L fermenter vessel. The maximum melanin concentration (Pmax) at 5L scale process reached 810.313+/-0.003 μg/mL at 120 h. Moreover, the overall melanin formation rate (Qp) of 0.88 μg/mL. Similarly, there observed an increase in cell biomass concentration with increase in time interval. The cell biomass reaches at its stationary concentration after 100h of incubation. The dissolved oxygen concentration (DO%) was also monitored from the control panel. There observed a sudden drop in DO% before 10 h of incubation. It shows that the organism is most active and they are at their exponential phase of growth during this time and so utilized more oxygen. The pH of the medium also showed a reduction with increase in time period.

**Figure 2.**
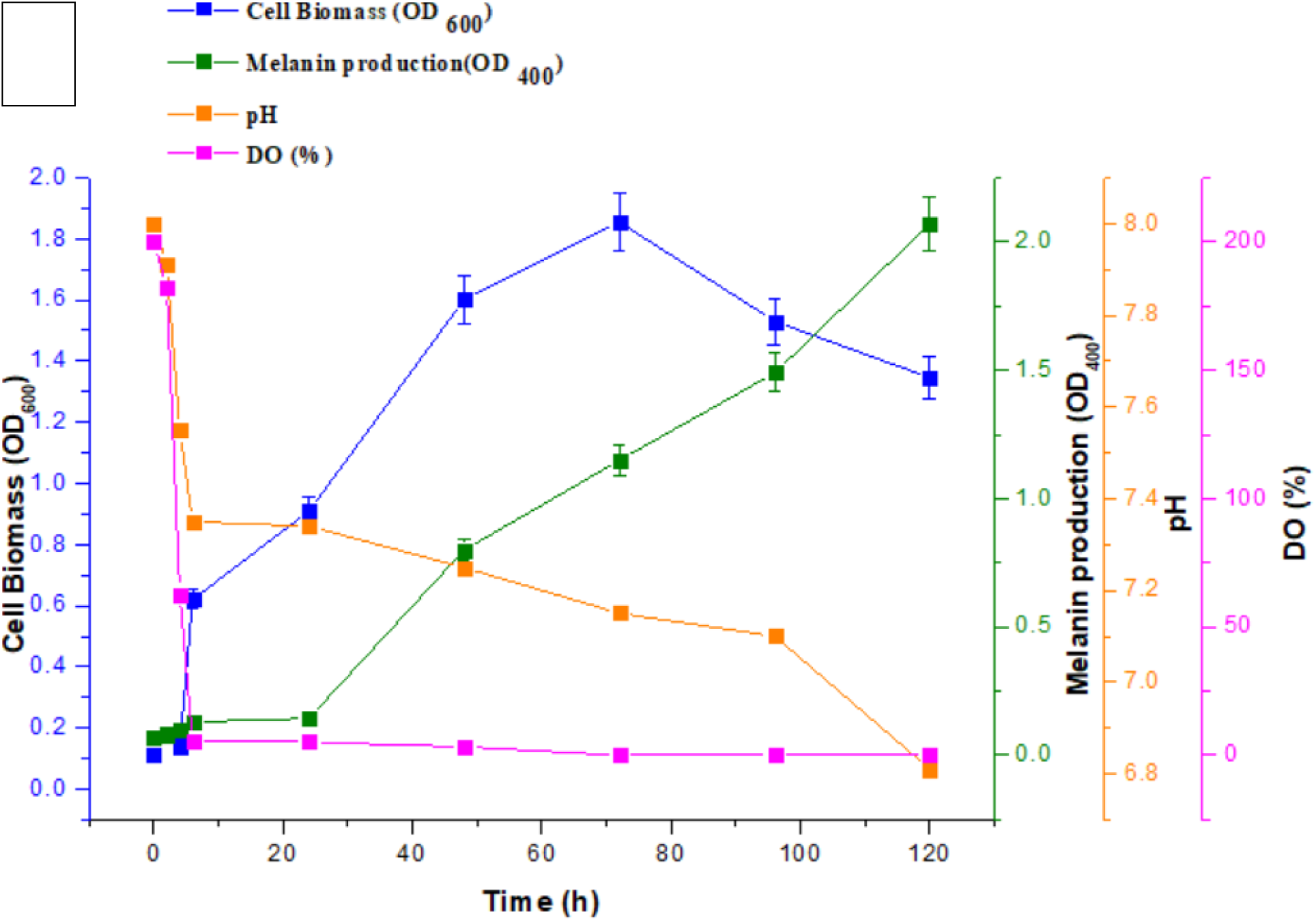
(a) Time dependent production of cell biomass, melanin, pH changes and dissolved oxygen concentration in 5L fermenter.

### 3.4 Melanin as nanoparticle

The nano structural properties of melanin were elucidated using different characterization method including UV spectral analysis, diffractive light scattering, zeta potential analysis, transmission electron microscopy etc. The melanin synthesized by *Pseudomonas stutzeri* BTCZ109 was found to be ∼5-7 nm size.

#### 3.4.1 Scanning electron microscope

Scanning electron microscope micrographs of the extracted bacterial melanin pigment produced by *Pseudomonas stutzeri* strain BTCZ109 at different magnification showed small clustered spheres lacking structural order (Fig 4(a)).

#### 3.4.2 UV-Visible Spectroscopy

In the present study (Fig 3(a,b)), the strain BTCZ109 showed an almost similar absorbance range with that of standard DOPA melanin and it was noted that bacterial strains melanin showed maximum absorbance at 213 nm.

**Fig 3:**
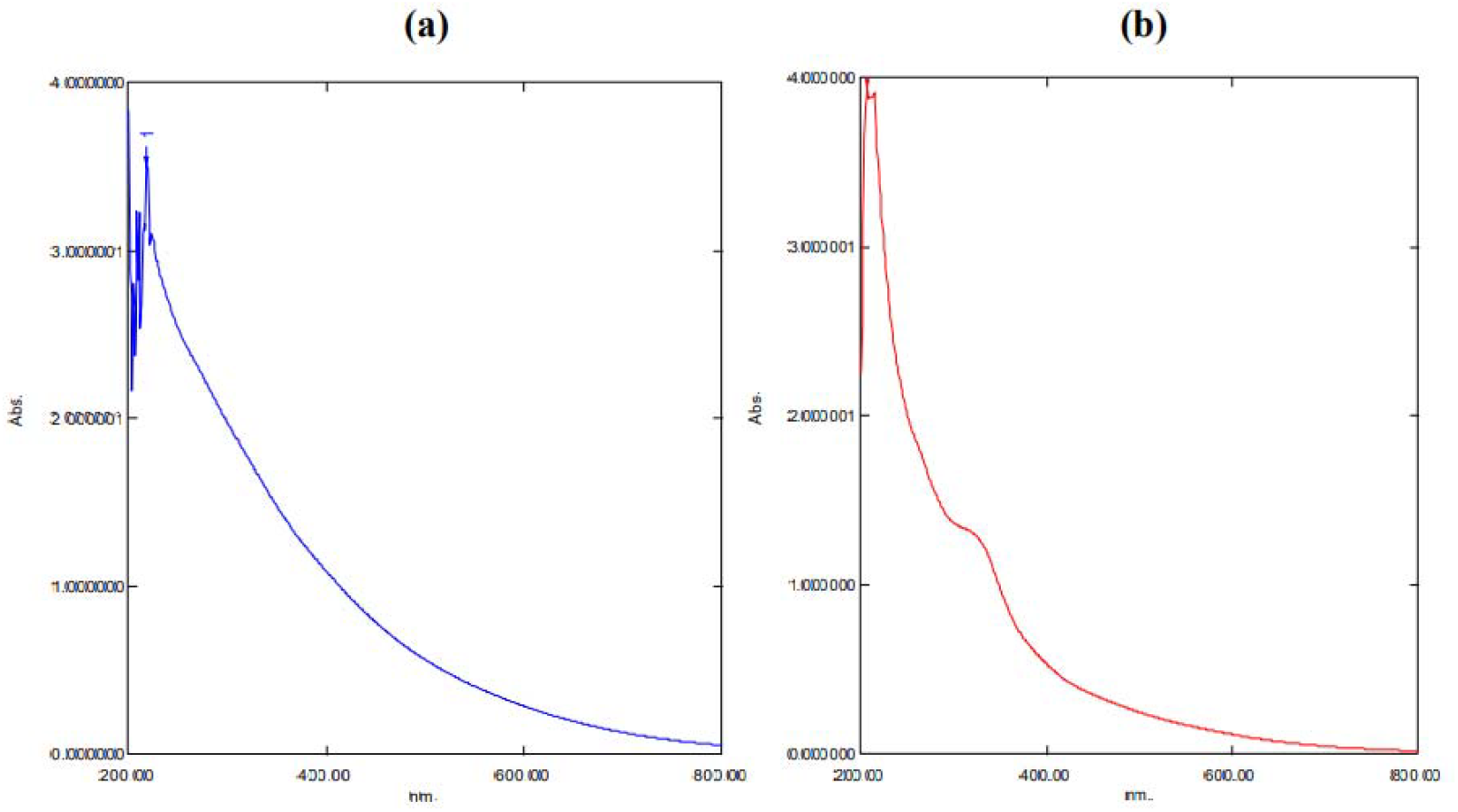
UV spectrum of melanin nanoparticle (a) DOPA melanin (b) bacterial melanin

#### 3.4.3 DLS Analysis

The melanin granules obtained after centrifugation and acid precipitation were sonicated and evaluated using diffractive light scattering, zeta potential and Transmission electron Microscopy to determine particle size distribution, stability and size range respectively.

It was inferred from the result that the -melanin granules synthesized by *Pseudomonas stutzeri* strain exhibited nano-sized distributions in the range of 107.07 nm (with a smaller intensity of 1% and width of 55.42) and 879.52 nm (with a smaller intensity of 7.29 % and width of 719) with a mean hydrodynamic diameter of 400.8 nm (Fig 4(f)). The increase in diameter was due to the existence of water molecules making non-covalent interactions with the melanin nano granules. Meanwhile, after sonication particles of size range ∼2-10 nm can be observed along with bigger sized particles with lesser hydrodynamic diameter. The zeta potential was -15.7 ±1.2 mV (Fig 4(g)). The negative zeta potential value indicated the electrostatic stability of this molecule in colloidal suspension. The increased stability with nano-sized distribution makes it an effective candidate in several therapeutic applications. Pyomelanin molecule with nano-sized distribution with potent anti-inflammatory and anti-cancer properties has already been reported (Narayanan *et al*., 2020).

**Fig 4:**
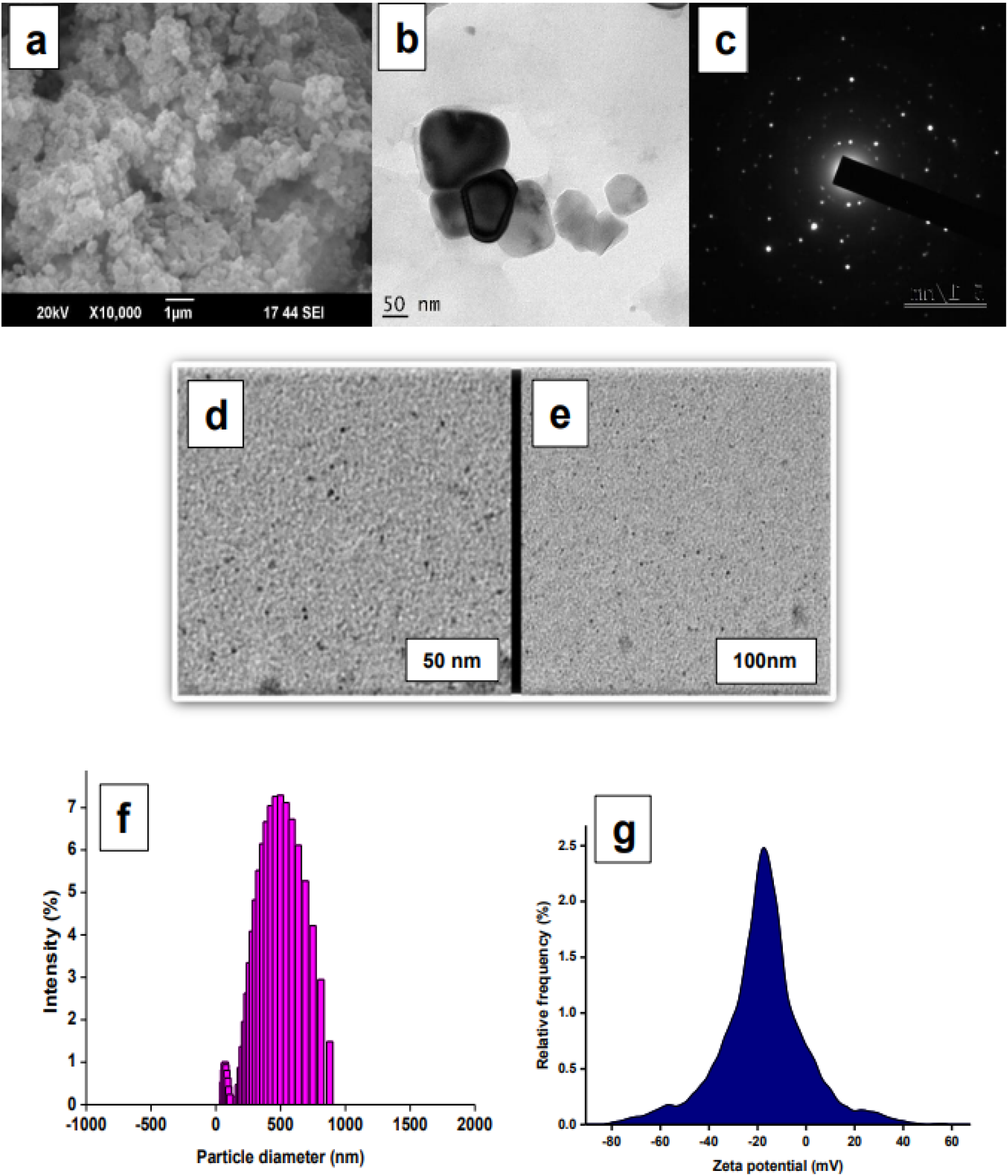
(a) SEM analysis (b) TEM image of melanin nanoparticle before sonication (c) SAED pattern and intensity profile (d) TEM images after sonication 50nm (e) 100nm (f) DLS Analysis (g) Zeta potential distribution

#### 3.4.4 TEM image

All the granules were almost spherical in geometry after sonication. Nano melanin granules under 50 nm were shown below. [Fig 4(b,c,d,e)]. The nano structural distribution of melanin particles was already reported in a previous study conducted in *Pseudoalteromonas* species (Narayanan *et al*., 2020).

## 4. Conclusion

Melanins are natural polymers synthesized by all forms of life. Here in this study large scale production of melanin from *Pseudomonas stutzeri* strain BTCZ109 was carried out in large scale fermenters. The factors optimized through One-factor-at-a-time method of optimization and statistical method of optimization was used while performing largescale production in fermenters. For this objective two different sized fermenters of 2L and 5L capacities were used. The L-tyrosine containing minimal media was used with all the optimized factors for this. 2L fermenter with 1L working capacity was used in which one impeller with four bladed turbines mounted on vertical shaft was there. The temperature was set at 37 ^0^C and at 140 rpm for 168 h time period to determine the maximum melanin production and cell biomass concentration. The cell biomass reached the maximum OD (Xmax) of 0.607 wet weight at 168 h, while the melanin concentration reached the maximum (Pmax) of 388.35 μg/mL at 168 h. The overall melanin formation rate (Qp) of 0.733 μg/mL l68 h. While in the case of 5L fermenter, 4L capacity was considered as working volume. a ratio of 1.5 height-to diameter and two vertically mount impellers with six-bladed turbines on a shaft was used for fermentation purpose. Water jacket connected externally in the fermenter vessel for maintaining the temperature. The main fermentation operating conditions were at temperature 37 ^0^C, aeration 0.5-1 l/p/m, agitation speed 140 rpm (2.3 Hz) and pH maintained at 8.0 using 2.0 mol of 1 L NaOH and 2.0 mol of 1L HCl. At this fermentation condition, the maximum melanin concentration (Pmax) at 5 L scale process reached 810.313+/-0.003 μg/mL at 120 h and the overall melanin formation rate (Qp) of 0.88 μg/mL. The melanin thus synthesized by the bacteria was found to be nano sized with the help of several characterization methods.

## 5. Abbreviations

DO: Dissolved Oxygen
%: Percentage
mM: MilliMolar
g/l: gram/litre
rpm: Rotations per minute
^0^C: degree Celsius

## 6. Acknowledgements

All authors acknowledge the support of DST FIST Phase I for financial support for fermenter purchase. The first author also acknowledges the support of Council of Scientific and Industrial Research (CSIR), Govt. of India for granting fellowships during the study (CSIR Award no: 09/239(0535)2018-EMR-1).

## 8. Author contribution

The first author did all the experimental work and wrote the draft of the manuscript under the supervision of her doctoral supervisor, and mentor, who modified the manuscript for publication.

## Reference

1. Ammanagi, A. I., Badiger, A. S., & Shivasharana, C. T. (2019). Bioprospecting of Fungi for Melanin fabrication: A Comprehensive. Int. J. Sci. Res. in Biological Sciences Vol, 6, 4.

2. Cao, W., Zhou, X., McCallum, N. C., Hu, Z., Ni, Q. Z., Kapoor, U., … & Gianneschi, N. C. (2021). Unraveling the structure and function of melanin through synthesis. Journal of the American Chemical Society, 143(7), 2622–2637. 10.1021/jacs.0c12322

3. E. Yabuuchi, A. Ohyama(1972) Characterization of “pyomelanin”-producing strains of Pseudomonas aeruginosa. International Journal of Systematic and Evolutionary Microbiology. 10.1099/00207713-22-2-53.

4. Guo, L., Li, W., Gu, Z., Wang, L., Guo, L., Ma, S., … & Chang, J. (2023). Recent Advances and Progress on Melanin: From Source to Application. International Journal of Molecular Sciences, 24(5), 4360. 10.3390/ijms24054360

5. Liu, R., Meng, X., Mo, C., Wei, X., & Ma, A. (2022). Melanin of fungi: From classification to application. World Journal of Microbiology and Biotechnology, 38(12), 228. 10.1007/s11274-022-03415-0

6. Michael, H. S. R., Subiramanian, S. R., Thyagarajan, D., Mohammed, N. B., Saravanakumar, V. K., Govindaraj, M., … & Ramesh Kumar, C. (2023). Melanin biopolymers from microbial world with future perspectives—A review. Archives of Microbiology, 205(9), 306. 10.1007/s00203-023-03642-5

7. Milne, N., Thomsen, P., Knudsen, N., Rubaszka, P., Kristensen, M., & Borodina, I. (2020). Metabolic engineering of Saccharomyces cerevisiae for the de novo production of psilocybin and related tryptamine derivatives. Metabolic engineering, 60, 25–36. 10.1016/j.ymben.2019.12.007

8. Neubauer, P. (2011). Towards faster bioprocess development. Biotechnology Journal, 6(8), 902–903. 10.1002/biot.201000413

9. Nielsen, J. (2013). Production of biopharmaceutical proteins by yeast: advances through metabolic engineering. Bioengineered, 4(4), 207–211. 10.4161/bioe.22856

10. Niyonzima, F. N. (2019). Production of microbial industrial enzymes. Acta Sci. Microbiol, 12, 75–89.

11. Park, J., Moon, H., & Hong, S. (2019). Recent advances in melanin-like nanomaterials in biomedical applications: a mini review. Biomaterials research, 23(1), 1–10. 10.1186/s40824-019-0175-9

12. Pralea, I. E., Moldovan, R. C., Petrache, A. M., IlieiJ, M., HegheiJ, S. C., Ielciu, I., … & Iuga, C. A. (2019). From extraction to advanced analytical methods: The challenges of melanin analysis. International journal of molecular sciences, 20(16), 3943. 10.3390/ijms20163943

13. Roy, S., & Rhim, J. W. (2022). New insight into melanin for food packaging and biotechnology applications. Critical Reviews in Food Science and Nutrition, 62(17), 4629–4655. 10.1080/10408398.2021.1878097

14. S. Narayanan, N.K. Kurian(2020) Ultra-small pyomelanin nanogranules abiotically derived from bacteria-secreted homogentisic acid show potential applications in inflammation and cancer. BioNanoScience. 10.1007/s12668-019-00689-x.

15. Sáez-Sáez, J., Wang, G., Marella, E. R., Sudarsan, S., Pastor, M. C., & Borodina, I. (2020). Engineering the oleaginous yeast Yarrowia lipolytica for high-level resveratrol production. Metabolic Engineering, 62, 51–61. 10.1016/j.ymben.2020.08.009

16. Sánchez, S., Chávez, A., Forero, A., García-Huante, Y., Romero, A., Sánchez, M., … & Ruiz, B. (2010). Carbon source regulation of antibiotic production. The Journal of antibiotics, 63(8), 442–459. 10.1038/ja.2010.78

17. Sauer, M., Porro, D., Mattanovich, D., & Branduardi, P. (2008). Microbial production of organic acids: expanding the markets. Trends in biotechnology, 26(2), 100–108. 10.1016/j.tibtech.2007.11.006

